# Loss of Bacterial Cell Pole Stabilization in *Caulobacter crescentus* Sensitizes to Outer Membrane Stress and Peptidoglycan-Directed Antibiotics

**DOI:** 10.1101/784066

**Authors:** Simon-Ulysse Vallet, Lykke Haastrup Hansen, Freja Cecillie Bistrup, Julien Bortoli Chapalay, Marc Chambon, Gerardo Turcatti, Patrick H. Viollier, Clare L. Kirkpatrick

## Abstract

Rod-shaped bacteria frequently localise proteins to one or both cell poles in order to regulate processes such as chromosome replication or polar organelle development. However, the role of such polar factors in responses to extracellular stimuli has been generally unexplored. We employed chemical-genetic screening to probe the interaction between one such factor from *Caulobacter crescentus*, TipN, and extracellular stress and found that TipN is required for normal tolerance of cell envelope-directed antibiotics, including vancomycin that does not normally inhibit growth of Gram-negative bacteria. Forward genetic screening for suppressors of vancomycin sensitivity in the absence of TipN revealed the TonB-dependent receptor ChvT as the mediator of vancomycin tolerance. Loss of ChvT improved resistance to vancomycin and cefixime in the otherwise sensitive Δ*tipN* strain. The activity of the two-component system regulating ChvT (ChvIG) was increased in Δ*tipN* cells relative to wild type under some, but not all, cell wall stress conditions that this strain was sensitised to, in particular cefixime and detergent exposure. Together, these results indicate that the ChvIG two-component system has been co-opted as a sensor of cell wall stress and that TipN can influence cell envelope stability and ChvIG-mediated signaling in addition to its roles in intracellular development.

**Author summary:** Maintenance of an intact cell envelope is essential for free-living bacteria to survive harsh conditions they may encounter in their environment. In the case of rod-shaped bacteria, the poles of the cell are potential weak points in the cell envelope due to the high curvature of the layers and the need to break and re-form parts of the cell envelope at the division plane in order to form new poles as the cells replicate and divide. We have found that TipN, a factor required for correct division and cell pole development in the rod-shaped bacterium, *Caulobacter crescentus*, is also needed for maintaining normal levels of resistance to cell wall-targeting antibiotics such as vancomycin and cefixime, which interfere with peptidoglycan synthesis. We also identified an outer membrane receptor, ChvT, that was responsible for allowing vancomycin access to the cells and found that the two-component system that negatively regulates ChvT production was activated by various kinds of cell wall stress. Presence or absence of TipN influenced how active this system was in the presence of cefixime or of the membrane-disrupting detergent sodium deoxycholate. Since TipN is normally located at the poles of the cell and at the division plane just before cells complete division, our results suggest that it is involved in stabilisation of these weak points of the cell envelope as well as its other roles inside the cell.

## Introduction

The asymmetrically dividing alpha-proteobacterium *Caulobacter crescentus* (hereafter *Caulobacter*) temporally and spatially regulates its cell cycle in order to propagate the correct positioning of the polar stalk and flagellum from one generation of cells to the next [1–4]. Among the regulatory factors that transmit positional information differentiating the cell poles from the rest of the cell body is the TipN polarity factor that localizes to the division plane and remains at the new pole once division is completed [5, 6]. This provides a molecular marker that defines the newest cell pole (ie. the one arising from the most recent cell division) and ensures that the flagellum assembly factors are subsequently recruited there. Cells lacking TipN have a flagellum placement defect, but also other deficiencies. For example, cells lacking TipN are defective in chromosome partitioning, likely due to loss of an interaction between TipN and ParA which is required for prompt and unidirectional segregation of the *parS*-ParB complex from one cell pole to the other after initiation of chromosome replication [7, 8], a phenotype unrelated to flagellum positioning. Therefore, TipN acts as a new-pole-specific multifunctional regulator that spatially integrates at least two vital cellular processes (cellular differentiation and chromosome partitioning).

If the correct function of TipN depends on its relocalization from flagellar pole to division plane at cytokinesis, how is this achieved? It is currently unknown what signal triggers delocalization of TipN away from the pole and its reassembly at the division septum, but it has been shown that TipN can interact with the septum-located Tol-Pal complex [9]. This complex regulates later stages of cell division and is required for connecting the inner membrane, peptidoglycan and outer membrane during the peptidoglycan remodeling that is necessary for septum formation between the two daughter cells. It is conceivable, although unproven, that the arrival of Tol-Pal at the division plane is the recruitment signal for TipN. Tol-Pal is required for outer membrane fission at cell division and remains at the newly divided cell pole (in complex with TipN) after separation of the daughter cells is complete. Unlike in *E. coli*, the *Caulobacter* Tol-Pal complex is essential [10] and it has been proposed that it is required for general cell envelope integrity in addition to carrying out the last stage of cell division [9]. Although TipN has not been shown to participate directly in maintenance of cell wall integrity, a genetic link between TipN and other cell envelope-located protein complexes (in addition to Tol-Pal) has been identified. We previously found that loss of function of TipN was synthetically lethal with the quinolone antibiotic nalidixic acid (Nal), and discovered that this was because Nal induces strong expression of an efflux pump of the RND family, named AcrAB2-NodT, while the normal target of Nal (DNA gyrase) was apparently unaffected, and that this overexpression was specifically intolerable to Δ*tipN* cells [11]. Deletion of the efflux pump improved, rather than decreased, Nal resistance, and seemed to be related to pump structure rather than function because a chemical inhibitor of efflux (PAβN, phenylalanine arginine β-naphthylamide) did not improve Nal resistance. However, the molecular mechanism by which TipN protects against the Nal-induced overexpression of this efflux pump is still not known.

As the outer membrane of Gram-negative bacteria is their first line of defence against toxic environmental factors, in bacteria that live in oligotrophic environments (such as *Caulobacter*) the outer membrane is equipped with systems to (a) detect and respond negatively to potential toxins, or (b) detect and respond positively to nutrients. *Caulobacter* encodes 105 putative signal transduction proteins and 65 TonB-dependent receptors in its genome [12], which are potentially important for environmental sensing and nutrient uptake, respectively. The TonB-dependent receptors have not been systematically characterized, but individual members of this family have been found to be specific receptors for vitamins or metal ions [13–16] or to be transcriptionally upregulated under conditions of carbon or nitrogen starvation [17]. Regulation of TonB-dependent receptors, as for outer membrane proteins in general, is often post-transcriptionally controlled by small noncoding RNAs (sRNAs) [18]. *Caulobacter* encodes 133 sRNAs [19], of which only a few have been characterized, but in these cases they appear to be components of stress-response signaling pathways [20, 21]. The sRNA ChvR was recently shown to downregulate a single TonB-dependent receptor of unknown function, ChvT, in response to acid stress, starvation or DNA damage [22]. ChvR transcription under these conditions was induced by the two-component system ChvIG, which is highly conserved among the alpha-proteobacteria and particularly important for the association of pathogenic [23, 24] or symbiotic [25] alpha-proteobacterial species with their hosts. However, in *Caulobacter* it seems to have been repurposed to sense other types of environmental stress, since this species does not associate with a host cell.

In this study we sought to clarify why high expression of an efflux pump should be deleterious to Δ*tipN* cells and whether it was possible to identify other environmental conditions which were similarly toxic to this strain, to pinpoint the molecular mechanism of TipN’s role in protection against AcrAB2-NodT overexpression. We show that the toxicity of the efflux pump overexpression arises from general intolerance of the Δ*tipN* strain to overexpression of this class of cell envelope-spanning systems and can also be elicited by heterologous overexpression of other efflux pump components. Moreover, by high-throughput chemical-genetic screening we found that this strain is also sensitized to cell wall targeting antibiotics, particularly vancomycin, and that the vancomycin sensitivity can be suppressed by loss of the TonB-dependent receptor ChvT. We observed activation of the ChvIG-induced promoter of ChvT’s regulatory RNA, *chvR*, during cell envelope stress conditions including efflux pump overexpression, presence of cell wall-targeted antibiotics and the detergent sodium deoxycholate. The Δ*tipN* strain was sensitised to detergent stress relative to the wild-type, and the detergent-induced activity of *chvR* promoter activity in Δ*tipN* cells was correspondingly higher. Overall, these findings indicate a previously unsuspected role for TipN in stabilization of the cell envelope of *Caulobacter* and that the presence or absence of TipN can influence cell wall stress signaling mediated by the ChvIG-ChvR-ChvT pathway.

## Results

### TipN is required for normal growth under conditions of cell envelope overload from AcrAB2-NodT overexpression

We previously identified an AcrAB-family multidrug efflux pump module, *acrAB2nodT*, that was strongly induced in response to Nal and whose induction was responsible for the Δ*tipN* mutant’s Nal sensitivity [11] (Fig 1A). However, at this stage we were lacking measurements of steady-state levels of pump components in basal and nalidixic acid-induced conditions and had not completely excluded the possibility that this strain was also sensitised to Nal-mediated but AcrAB2-NodT-independent toxicity. We therefore carried out western blotting against AcrA to assess basal and Nal-induced protein levels of this component of the pump, as well as the levels obtained from induction of *acrA* gene expression from the vanillate-inducible promoter in the constructs we had previously used to investigate whether induced expression of *acrA* or *acrB2* independent of Nal could exert the same effect. However, we found that the expression from this construct was so low that it did not approach the basal level of AcrA expression from its native promoter, let alone the Nal-induced expression level (Fig 1B and S1 Fig), explaining why we had previously seen no phenotype using these constructs [11]. We therefore made alternative constructs using a vector carrying the xylose-inducible promoter upstream of the individual *acrA, acrB2* or *nodT* genes, *acrAB2*, or the whole operon. With this construct, expression levels in the absence of xylose (inducer) or the presence of glucose (repressor) were similar to the basal level of expression from the native promoter, while xylose-induced expression from this construct was comparable to Nal-induced expression from the native promoter (Fig 1C). In the Δ*tipN* mutant strain, xylose-induced overexpression of *acrA*, *acrAB2* or *acrAB2nodT* inhibited growth of the cells compared to the glucose-treated control (Fig 1D) while the wild type (WT) showed some partial sensitivity to overexpression of the whole operon and *acrAB2* genes but no effect of the *acrA* or *acrB2* genes alone (Fig 1E). The decreased growth seen in these end-point measurements corresponded to the cultures entering stationary phase at a lower OD600 while the growth rate in exponential phase was not affected (S2 Fig). These results suggest that the sensitivity of the Δ*tipN* mutant to Nal is independent of non-specific Nal-mediated toxicity and show that Δ*tipN* is more sensitive than WT to overproduction of individual components of the pump, not only the whole operon, suggesting that it is susceptible to non-specific overload of the cell envelope unrelated to drug efflux activity.

**Fig 1.**
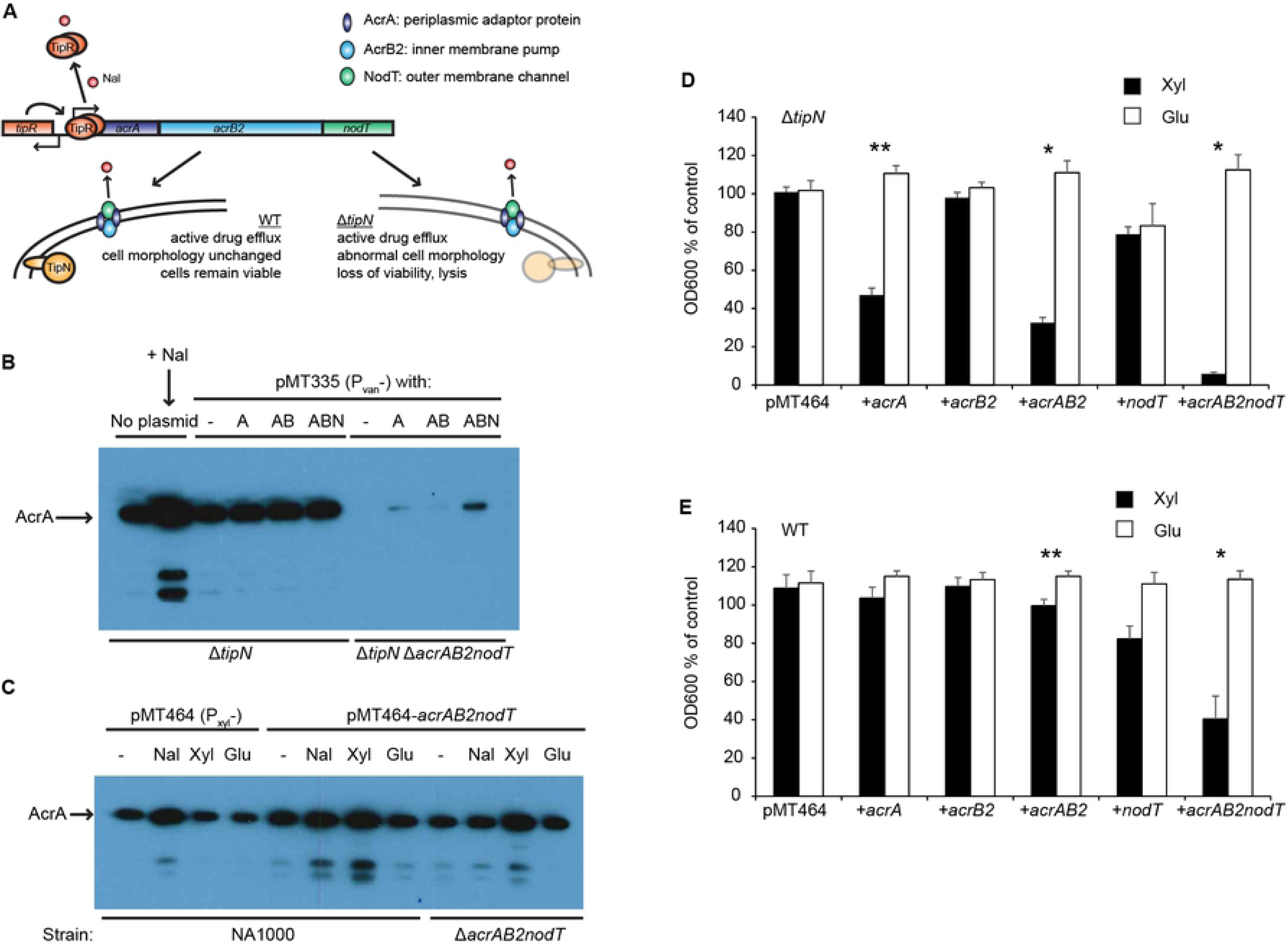
Overexpression *in trans* of *acrAB2nodT* efflux pump components precisely mimics the Nal sensitivity phenotype in Δ*tipN* cells. (A) Graphical model of the regulatory effect of nalidixic acid (Nal) on the *acrAB2nodT* operon and its consequences in WT and Δ*tipN* strains. Nalidixic acid causes the repressor TipR to leave the promoter, resulting in high expression of the efflux pump genes and their incorporation into the cell envelope. (B) Immunoblotting with specific antibody against AcrA of cell extracts of Δ*tipN* and Δ*tipN* Δ*acrAB2nodT* with no plasmid or with pMT335-based constructs in which expression of the cloned genes (-, empty vector; A, *acrA*; AB, *acrAB2*; ABN, *acrAB2nodT*) was induced from the vanillate-inducible promoter with 50 μM vanillate for 3hr before cell harvesting. Δ*tipN* cells without the plasmid were treated with 20 μg/ml nalidixic acid for 3hr before harvesting. Cell cultures were normalised by OD600 before harvesting to ensure equivalent loading in all lanes of the gel. This image was deliberately overexposed to allow for visualisation of the weakest bands; a less exposed image of the same membrane is provided in S1 Fig. (C) Immunoblotting with specific antibody against AcrA of cell extracts of NA1000 (WT) or Δ*acrAB2nodT* containing the empty vector pMT464 or the pMT464-*acrAB2nodT* overexpression vector in which expression of the cloned genes is induced with 0.3% xylose (or repressed by 0.2% glucose) for 3hr before cell harvesting. The same strains were also treated with 20 μg/ml nalidixic acid for 3hr before harvesting in order to directly compare nalidixic acid induction from the native promoter to xylose-induced overexpression from the pMT464 promoter in the same gel. Cell cultures were normalised by OD600 before harvesting to ensure equivalent loading in all lanes of the gel. (D) Growth assay of Δ*tipN* cells with empty vector pMT464 and the series of pMT464 overexpression plasmids containing *acrAB2nodT* component genes separately or together, treated with 0.3% xylose or 0.2% glucose for 20hr before culture density measurement. Data are expressed as % of OD600 of control cultures of each strain in untreated PYE. (E) Growth assay of WT cells with empty vector pMT464 and the series of pMT464 overexpression plasmids containing *acrAB2nodT* component genes separately or together, treated with 0.3% xylose or 0.2% glucose for 20hr before culture density measurement. Data are expressed as % of OD600 of control cultures of each strain in untreated PYE. Statistical significance is denoted by * for p < 0.05 and ** for p < 0.01 in pairwise (Xyl. vs. Glu) comparisons.

### AcrAB2-NodT is not intrinsically toxic

To test whether this toxicity was unique to *acrAB2nodT* or if it could be induced by other efflux pumps, we cloned the *Caulobacter acrA3* and *acrAB3* genes, the *E. coli acrA* and *acrAB* genes and the *mexA* and *mexAB* genes of the *Pseudomonas aeruginosa* multidrug efflux pump MexAB-OprM in order to carry out heterologous overexpression growth assays. Consistent with a non-specific effect, we observed that the Δ*tipN* mutant was sensitised to overexpression of the *acrA3* (*Caulobacter*) and *mexA* (*P. aeruginosa*) genes relative to WT (Fig 2A). The WT strain was also sensitised to overexpression of the *acrAB3*, *E. coli acrAB* and *Pseudomonas mexAB* gene pairs (Fig 2B), suggesting that heterologous overexpression of foreign (or endogenous but not usually strongly expressed) complexes of periplasmic adaptor together with inner membrane pump components is not well tolerated in *Caulobacter*. This genetic approach is limited in that the relative expression level and stability of the overexpressed proteins is unknown and may theoretically differ between constructs, but the increased sensitivity of the Δ*tipN* strain to *acrA3* and the *Pseudomonas* homologue *mexA* compared to WT still supports the hypothesis that the toxicity arises from a general overexpression sensitivity of the Δ*tipN* cells and not of any specific intrinsic toxicity of the AcrAB2-NodT complex.

**Fig 2.**
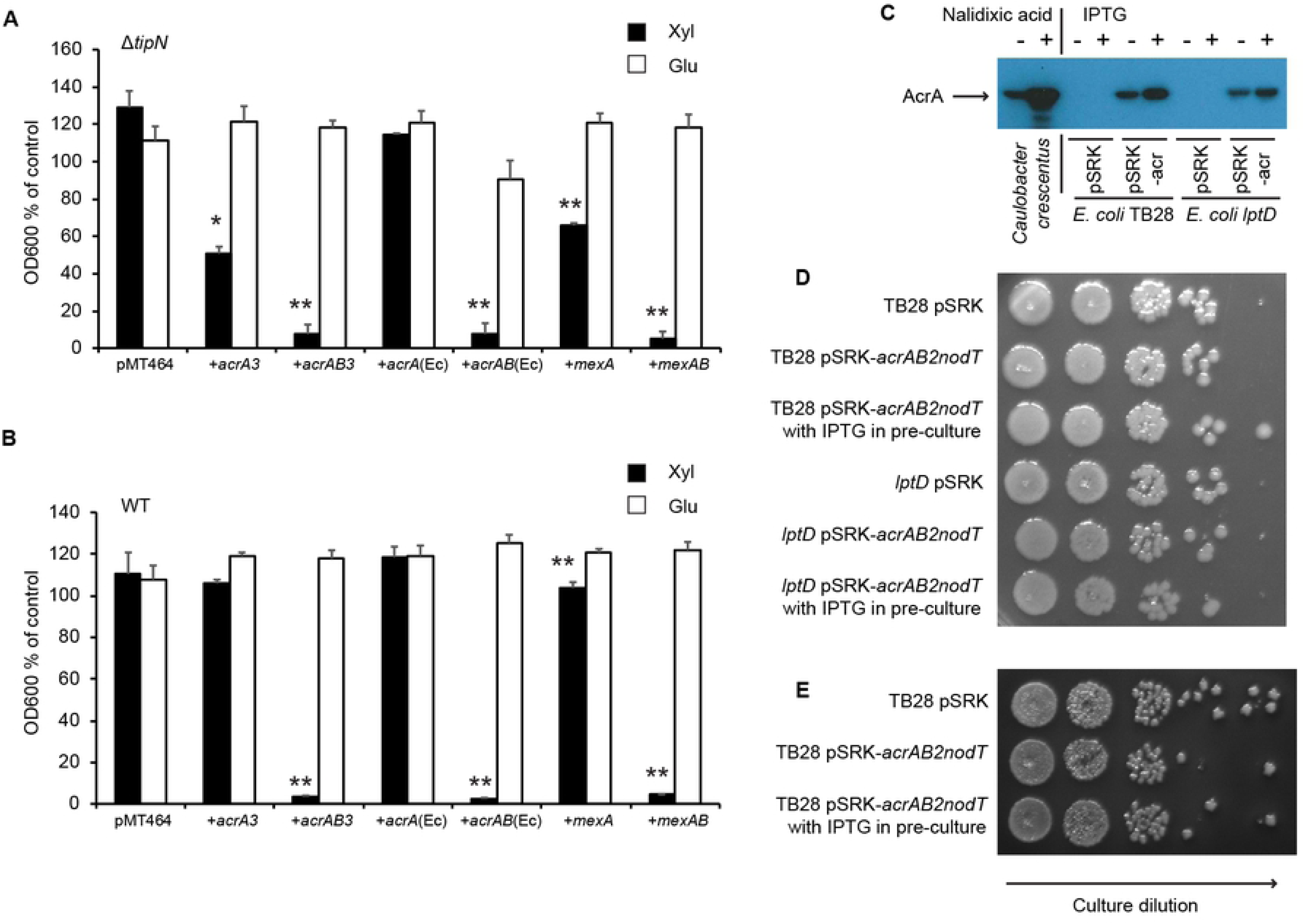
Heterologous overexpression of efflux pump components affects *Caulobacter* WT and Δ*tipN*, and *E. coli* WT and *lptD*, differently. (A) Growth assay of Δ*tipN* cells with empty vector pMT464 and the heterologous overexpression constructs for induced overexpression of *Caulobacter acrA(B)3*, *E. coli acrA(B)* and *P. aeruginosa mexA(B)*, treated with 0.3% xylose or 0.2% glucose for 20hr before culture density measurement. Data are expressed as % of OD600 of control cultures of each strain in untreated PYE. (B) Growth assay of WT cells with empty vector pMT464 and the heterologous overexpression constructs for induced overexpression of *Caulobacter acrA(B)3*, *E. coli acrA(B)* and *P. aeruginosa mexA(B)*, treated with 0.3% xylose or 0.2% glucose for 20hr before culture density measurement. Data are expressed as % of OD600 of control cultures of each strain in untreated PYE. Statistical significance is denoted by * for p < 0.05 and ** for p < 0.01 in pairwise (Xyl. vs. Glu) comparisons. (C) Immunoblotting with specific antibody against AcrA in cell extracts prepared from *Caulobacter* WT treated or not with 20 μg/ml nalidixic acid for 3hr before harvesting, alongside *E. coli* cell extracts containing empty vector pSRK-Km or the overexpression construct pSRK-*acrAB2nodT*, with or without 1 mM IPTG for 3hr before harvesting. Cell cultures were normalised by OD600 before harvesting to ensure equivalent loading in all lanes of the gel. (D) Viability assay of WT *E. coli* (TB28) and *lptD* mutant with empty vector pSRK-Km or the overexpression construct pSRK-*acrAB2nodT* with or without 1 mM IPTG in the pre-culture, on LB agar with 1 mM IPTG. Starter cultures were diluted into LB with or without 1 mM IPTG and grown to exponential phase before harvesting, normalising by OD600, serially tenfold diluting and plating 5 μl spots of the diluted cells. Representative image of two biological replicates. (E) Viability assay of WT *E. coli* (TB28) with empty vector pSRK-Km or the overexpression construct pSRK-*acrAB2nodT* with or without 1 mM IPTG in the pre-culture, on McConkey agar containing glucose as carbon source and 1 mM IPTG. Starter cultures were diluted into LB with or without 1 mM IPTG and grown to exponential phase before harvesting, normalising by OD600, serially tenfold diluting and plating 5 μl spots of the diluted cells. Representative image of two biological replicates.

We also performed the reverse heterologous overexpression experiment of cloning the *Caulobacter acrAB2nodT* operon into a vector in which we could overexpress it in *E. coli*, and tested its effect on growth in a *lac*^-^ but otherwise wild type *E. coli* strain (TB28) and an *E. coli lptD* mutant that has a destabilised outer membrane (allele formerly known as *imp4213*, [26]). Basal expression of AcrA from this vector was similar to the basal level in *Caulobacter*, while IPTG-induced expression was stronger and similar in both *E. coli* strains, although it was less strong than Nal-induced expression from the *Caulobacter* native promoter (Fig 2C). However, no toxic effect of *acrAB2nodT* overexpression was seen in either strain growing on LB (Fig 2D), nor for the TB28 strain growing on McConkey agar, on which the *lptD* mutant cannot grow (Fig 2E). Hence, *acrAB2nodT* overexpression is not toxic in a heterologous overexpression system, even under genetic or environmental membrane-destabilising conditions, suggesting that its effect on the *Caulobacter* Δ*tipN* mutant is due to this strain’s intolerance of cell envelope protein complex overexpression.

### TipN is required for maintenance of normal cell morphology and division under AcrAB2-NodT-mediated cell envelope stress

To investigate how efflux pump overexpression could be exerting its toxic effect, we studied cell morphology of WT and Δ*tipN* strains overexpressing the efflux pump (whole operon or individual components) from the xylose-inducible promoter. Compared to the empty vector control, Δ*tipN* cells overexpressing the *acrA* gene became filamentous, while overexpressing the *acrAB2* genes in Δ*tipN* caused bulging, branching and frequent lysis in addition to filamentation (Fig 3A), similar to Δ*tipN* cells exposed to Nal (Fig 3B). WT cells had mild bulging and filamentation defects with Nal and with *acrAB2nodT* operon overexpression, while overexpression of the *acrA* or *acrAB2* constructs alone had no effect. To confirm that the Nal-induced cell morphology defect was due to efflux pump overexpression from its native promoter, we imaged identically treated Δ*tipN* and Δ*tipN* Δ*acrAB2nodT* cells, and as expected, the deletion of the *acrAB2nodT* pump genes protected Δ*tipN* cells from the Nal-induced morphology defect (Fig 3C). Together, these experiments suggest that TipN is required for maintenance of correct cell division during cell envelope overload with periplasmic proteins or complexes.

**Fig 3.**
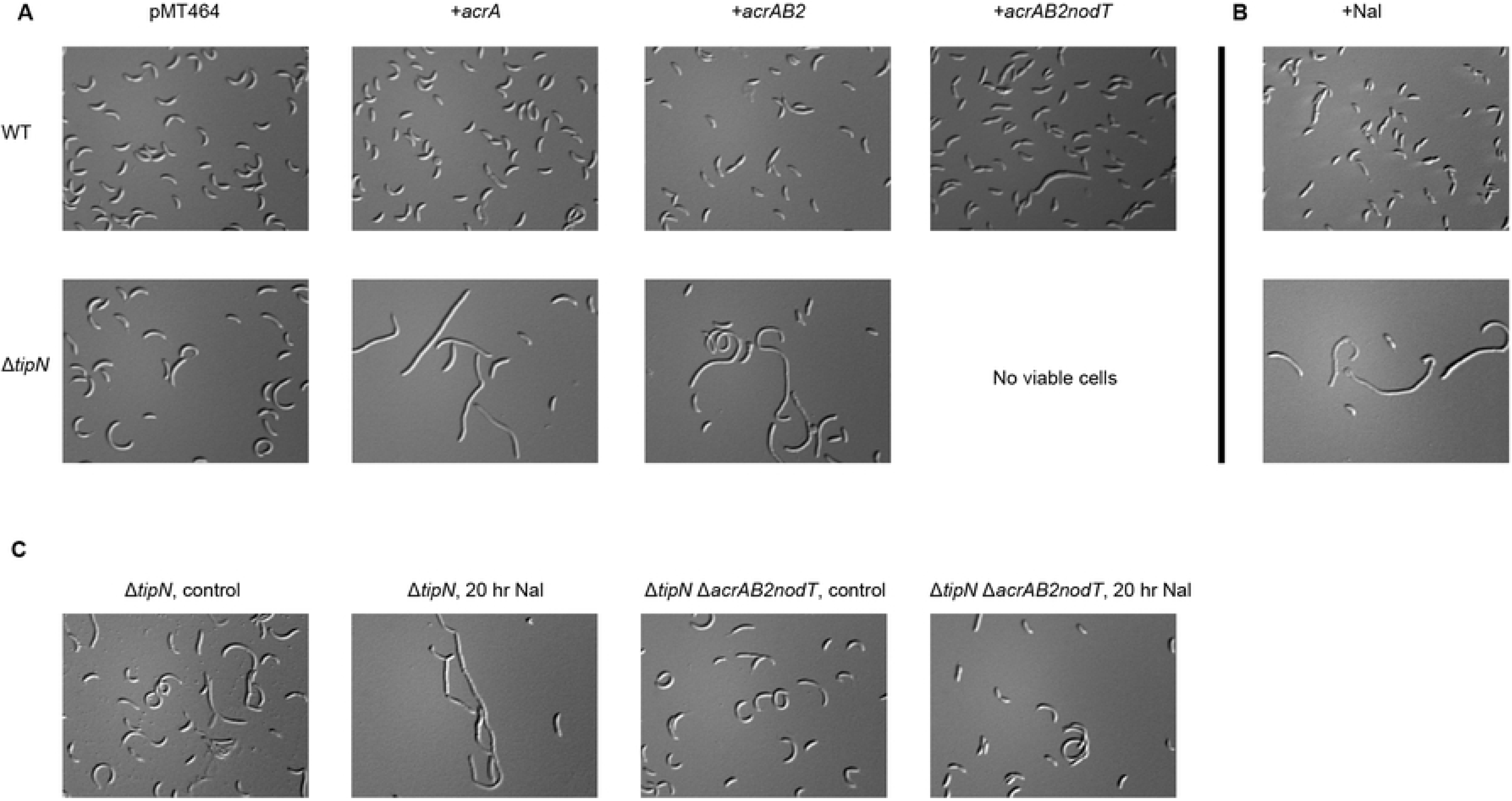
Nal exposure and *acrAB2nodT* overexpression cause identical cell morphology defects in Δ*tipN* cells. (A) Differential interference contrast (DIC) images of WT and Δ*tipN* cells containing pMT464, pMT464-*acrA*, pMT464-*acrAB2* or pMT464-*acrAB2nodT* and treated with 0.3% xylose for 20hr before imaging. Control cultures of both strains treated with 0.2% glucose were prepared and imaged in every biological replicate and had identical cell morphology to the empty vector control cultures. (B) DIC images of WT and Δ*tipN* cells treated with 20 μg/ml Nal for 20hr before imaging. (C) DIC images of Δ*tipN* and Δ*tipN* Δ*acrAB2nodT* cells treated or not with 20 μg/ml Nal for 20hr before imaging.

### Chemical genetic screening reveals that Δ*tipN* cells are also sensitised to cell wall-targeting antibiotics independent of AcrAB2-NodT

Having established that the Δ*tipN* mutant is susceptible to cell wall stress caused by overload of cell envelope located proteins, specifically efflux pump components, we then investigated whether we could identify any sensitivity to small molecule stressors that mimicked this effect by chemical genetic screening. We screened WT *Caulobacter* against the Prestwick Chemical Library in the presence and absence of Nal, reasoning that any drugs which could inhibit growth in the presence of Nal but not in its absence, would be synthetically toxic with Nal-induced AcrAB2-NodT overexpression. They would therefore act as a chemical phenocopy of the Δ*tipN* mutant phenotype, in which case their mechanism(s) of action could indicate why TipN is protective against cell wall stress as described above. After identifying the compounds that fitted this criterion, we compared the list of hits from this screen against those that had been identified as Δ*tipN*-specific growth inhibitors in a previous study [27]. This was done in order to differentiate between hits that were synergistic with Nal but unrelated to its activity as transcriptional inducer of AcrAB2-NodT expression, from hits which were likely to be synergistic with Nal-induced AcrAB2-NodT expression, to minimise false positives and avoid analysing irrelevant phenotypes. Strikingly, the compound which showed the strongest Δ*tipN*-dependent and Nal-dependent growth inhibition was vancomycin (Vanco) (Fig 4A). We also identified the 3^rd^-generation cephalosporin cefixime as a hit (albeit low-scoring) which fitted these criteria, and noted that a structurally similar drug, cefotaxime, had been a low-scoring hit in the Δ*tipN-*specific screen although it did not show Nal-dependent growth inhibition in WT (S3 Fig). Together, these results suggested that the Δ*tipN* mutation could cause cell envelope instability that allows access of cell wall-targeting drugs to the periplasm and sites of peptidoglycan synthesis. To validate the screening data, we performed growth assays to test the dose dependence of Vanco sensitivity in WT and Δ*tipN* cells and whether the Δ*acrAB2nodT* deletion could protect Δ*tipN* cells from Vanco as it did from Nal. Consistent with the screening experiment, the Δ*tipN* mutant was sensitised to Vanco relative to WT at 12.5 μg/ml (in the high-throughput screen Vanco was present at 10 μM = 15 μg/ml) (Fig 4B) and the addition of Nal sensitised all strains still further (Fig 4C). Decreased end-point growth measurements corresponded to entry into stationary phase at a lower OD600 (S5 Fig). However, the Δ*acrAB2nodT* deletion did not protect the WT or Δ*tipN* strains against the synergistic effect of Nal + Vanco, which would have been expected if Vanco had acted as a true phenocopy of the Δ*tipN* mutant phenotype at the genetic as well as phenotypic level. Therefore, although the presence of Nal, and presumably the subsequent *acrAB2nodT* overexpression, had been sufficient to cause increased Vanco sensitivity in WT cells, the Vanco sensitivity was not fully dependent on *acrAB2nodT* as it had been for the Δ*tipN* mutant’s Nal sensitivity. Imaging of WT and Δ*tipN* cells after 24 hr Vanco treatment showed that WT cells were not affected at the level of cell morphology, while the Δ*tipN* strain showed frequent signs of lysis, especially blebbing at the cell poles and division plane (where TipN is normally located in WT cells) which we had only infrequently observed in Nal-treated Δ*tipN* cells (Fig 4D, compare to Fig 3B and 3C). Hence, although the chemical-genetic screening results had initially indicated that the Nal and Vanco toxicity to the Δ*tipN* strain were linked, our follow-up validation experiments have shown that Nal and Vanco exert distinct cell envelope-directed toxic effects in the absence of TipN, presumably through independent pathways.

**Fig 4.**
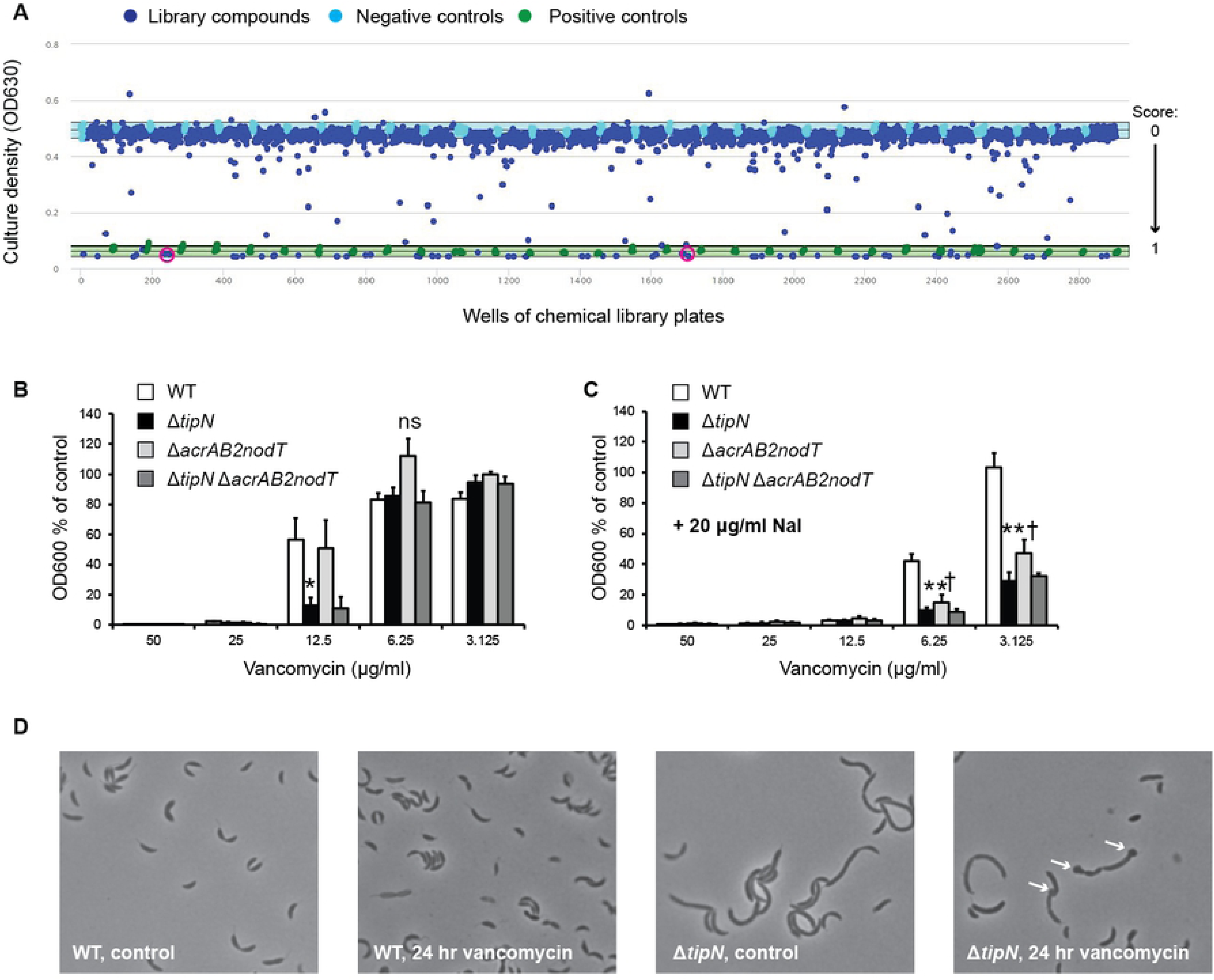
Nal and Vanco are synergistically lethal. (A) Scatter plot of the results of screening WT *Caulobacter* against the Prestwick Chemical Library for Nal-dependent growth inhibitors (all library compounds at 10 μM and Nal at 20 μg/ml in all conditions including the controls). The vertical axis indicates culture growth (measured by OD630) and the horizontal axis denotes the number of wells. Blue points correspond to wells containing library compounds, green points are the positive control wells (2 μg/ml=6.25 μM chloramphenicol) and light blue points are negative controls containing DMSO at 0.1% (final concentration of the diluent of the library compounds; does not inhibit growth). The light blue band behind the negative control points corresponds to mean ± 3 standard deviations around the mean of the negative controls. The Z’ score for this screen was 0.894 and the Z’ score for the corresponding control screen, of WT cells without Nal, was 0.820. Pink circles denote the wells containing Vanco. Vanco had a similarly close score to those of the positive controls in the Δ*tipN* screen (0.88) as it did in the Nal dependence screen shown here (1.00). (B) Growth assay of WT, Δ*tipN*, Δ*acrAB2nodT* and Δ*tipN* Δ*acrAB2nodT* cultures treated with Vanco at the indicated concentrations for 20hr before culture density measurement. Data are expressed as % of OD600 of control cultures of each strain in untreated PYE. (C) Growth assay of WT, Δ*tipN*, Δ*acrAB2nodT* and Δ*tipN* Δ*acrAB2nodT* cultures treated with Vanco at the indicated concentrations and 20 μg/ml Nal for 20hr before culture density measurement. Data are expressed as % of OD600 of control cultures of each strain in untreated PYE. Statistical significance is denoted by: * for p < 0.05 and ** for p < 0.01 in WT to Δ*tipN* comparison and † for p < 0.05 in WT to Δ*acrAB2nodT* comparison (“ns” signifies “not statistically significant”). (D) Phase-contrast images of WT and Δ*tipN* cells treated or not with 15 μg/ml Vanco for 24hr before imaging. White arrows indicate bleb formation sites at poles and division plane of cells.

### Vancomycin sensitivity of Δ*tipN* is suppressed by loss of the TonB-dependent outer membrane receptor ChvT

To search for genetic suppressors of the Δ*tipN* Vanco sensitivity phenotype, we performed forward genetic screening by constructing a pooled transposon mutant library in the Δ*tipN* strain and enriching it in Vanco-containing medium for individual clones that were capable of forming single colonies on media with 15 μg/ml Vanco. From this screen, we mapped three independent transposon insertions in or near the *CCNA_03108* gene (Fig 5A), annotated as a TonB-dependent outer membrane receptor of unknown substrate specificity named ChvT [22]. Two transposon insertions (2v15 and 3l15) mapped to the coding sequence, while one (1h15) appeared to be located in the putative promoter. Cloning of the *chvT* promoter region from the WT strain and the 1h15 transposon mutant into a β-galactosidase transcriptional reporter showed that (1) the 1h15 insertion in P*_chvT_*abolished transcriptional activity, (2) the promoter was active in both Δ*tipN* and WT strains and (3) that it did not respond to the presence either of Nal or Vanco (Fig 5B). Therefore, the promoter insertion is likely a loss of function allele.

**Fig 5.**
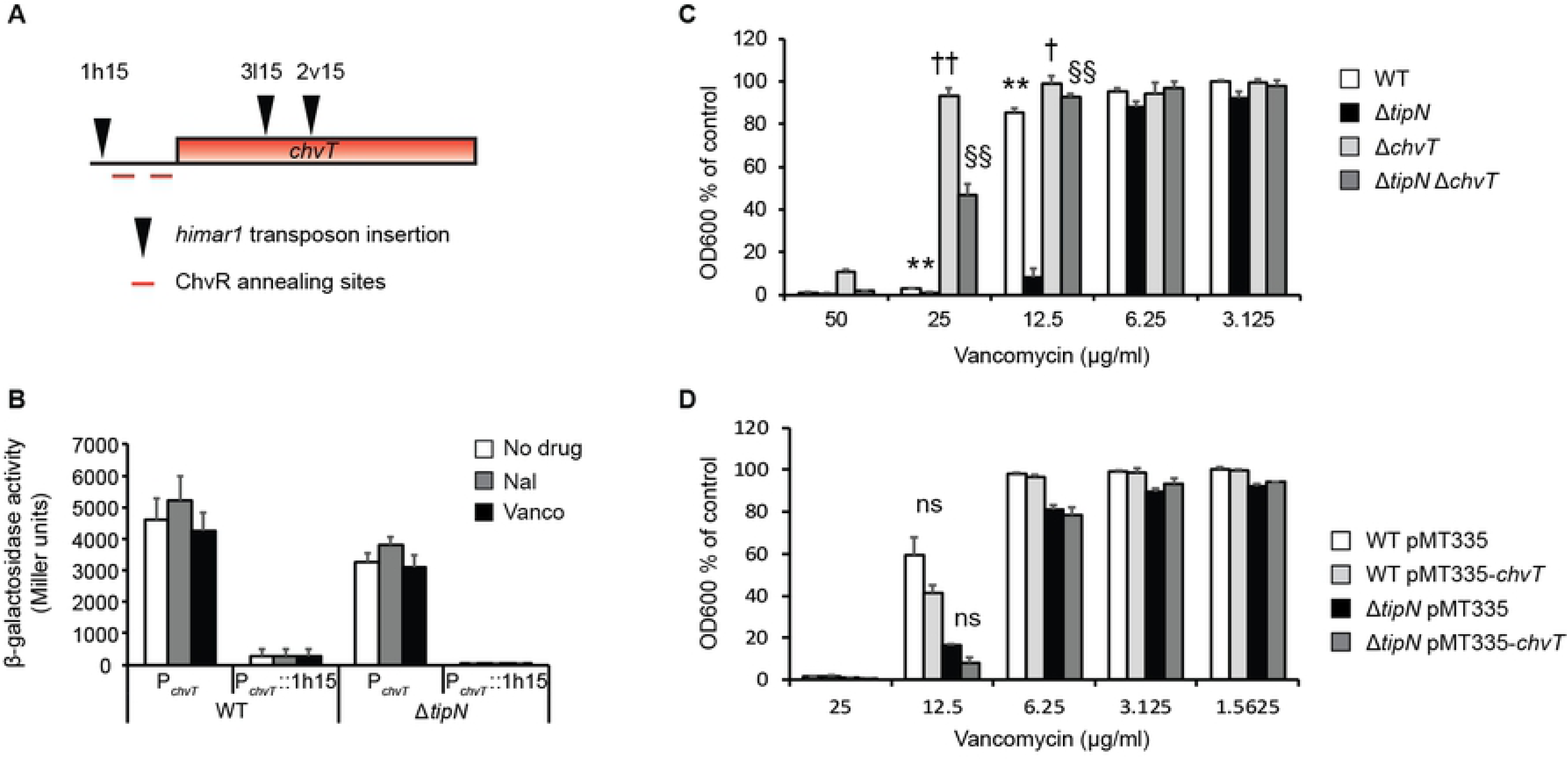
Loss of ChvT confers Vanco resistance. (A) Positions (approximately to scale) of the 1h15, 2v15 and 3l15 *himar1* transposon insertions in the *chvT* coding region or promoter that conferred Vanco resistance to the Δ*tipN* mutant. (B) Promoter activity of P*_chvT_*(WT promoter or mutant promoter containing the 1h15 *himar1* transposon insertion) in WT and Δ*tipN* strains in untreated PYE medium, 20 μg/ml Nal or 15 μg/ml Vanco, for 3hr. (C) Growth assay of WT, Δ*tipN*, Δ*chvT* and Δ*tipN* Δ*chvT* treated with Vanco at the indicated concentrations for 20 hr before culture density measurement. Data are expressed as % of OD600 of control cultures of each strain in untreated PYE. Statistical significance is denoted by: ** for p < 0.01 in WT to Δ*tipN* comparison, † for p < 0.05 and †† for p < 0.01 in WT to Δ*chvT* comparison and §§ for p < 0.01 in Δ*tipN* to Δ*tipN* Δ*chvT* comparison. (D) Growth assay of WT and Δ*tipN* containing the overexpression plasmid pMT335-*chvT* or the pMT335 empty vector, treated with 50 μM vanillate and Vanco at the indicated concentrations for 20hr before culture density measurement. Data are expressed as % of OD600 of control cultures of each strain in PYE containing 50 μM vanillate only (“ns” signifies “not statistically significant”).

In support of our hypotheses that (1) loss of function of *chvT* protects against Vanco and (2) the Vanco and Nal sensitivity phenotypes are unlinked, the double deletion mutant Δ*tipN* Δ*chvT* grew better in Vanco similarly to the 3l15 transposon insertion, but neither of these was protected against Nal, and the combination of Nal and Vanco was still highly toxic to all strains suggesting that the two antibiotics can exert a synergistic effect that Δ*chvT* cannot defend against (S4 Fig). The Δ*chvT* deletion also protected WT *Caulobacter* against high concentrations of Vanco, showing that it is not specifically associated with the Δ*tipN* genotype (Fig 5C). However, overexpression of *chvT* did not cause any significant sensitivity to Vanco in either WT or Δ*tipN* (Fig 5D), indicating that it is the presence or absence of ChvT that dictates the phenotype and that upregulating ChvT does not influence it. We performed the same experiments with cefixime and cefotaxime and confirmed that the Δ*tipN* mutant is sensitised to both of these antibiotics, but found that the Δ*chvT* deletion was less effective in protection against cefixime and cefotaxime than it had been for Vanco and *chvT* overexpression had no effect on growth in the presence of these antibiotics (S6 Fig). Finally, we tested whether Δ*chvT* could protect the Δ*tipN* mutant against overexpression of *acrAB2nodT* from the xylose-inducible promoter and, consistent with its lack of effect against Nal, we did not observe any change relative to the Δ*tipN* single mutant (S4 Fig; compare to Figure 1D). Therefore, loss of ChvT protects Δ*tipN* against cell wall stress when induced by the cell wall targeting antibiotics vancomycin, and to a lesser extent against cefixime and cefotaxime, but not against cell wall stress caused by other factors.

### Δ*tipN* displays elevated ChvIG-ChvR-ChvT signalling activity under outer membrane, but not periplasmic, stress conditions

It was recently shown that ChvT is the sole target of the sRNA ChvR, which binds to the 5’ UTR of the *chvT* mRNA and downregulates ChvT production [22]. Since loss of ChvT restored Vanco resistance to the Δ*tipN* strain, we hypothesised that the signalling pathway leading to transcription of ChvR, which is mediated by the two-component system ChvIG, could be altered in Δ*tipN* cells. Since it is known to respond to external factors such as starvation and acid stress, both in *Caulobacter* [22] and in other alpha-proteobacteria in which ChvIG is conserved [28], we investigated whether it also responds to the cell wall-targeting drugs identified in our chemical screen or to other sources of cell wall stress. We constructed a transcriptional P*_chvR_*-*lacZ* reporter and tested its activity in WT, Δ*tipN*, Δ*chvT*, Δ*tipN* Δ*chvT* and Δ*chvIG-hprK* strains (this last acts as a negative control for P*_chvR_*activity, as the two-component system ChvIG is the only known regulator of *chvR* transcription and therefore should display no or minimal P*_chvR_*activity). We measured the activity of this promoter in exponential and stationary phase in rich (PYE) and minimal (M2G) media to assess whether the promoter activity of this construct increased under conditions which upregulated ChvR RNA levels. Consistent with existing data [22] there was very low promoter activity in exponential phase in PYE, while it became active in M2G (Fig 6A). However, we also observed increased promoter activity in stationary phase in PYE as well as in M2G (Fig 6B), indicating that this promoter starts to become active at later stages of growth even in rich medium. Having validated that our reporter responded as expected to the M2G control condition, we tested whether it was also affected by overexpression of *acrAB2nodT*. Xylose-induced *acrAB2nodT* overexpression caused increased activity of P*_chvR_*compared to the glucose or empty vector controls in both WT and Δ*tipN* strains (Fig 6C), showing that the ChvIG-*chvR* signalling pathway can respond to loading of the cell envelope with AcrAB2-NodT efflux pump complexes. We then investigated the effect of Vanco, cefixime and cefotaxime on the P*_chvR_* promoter to see if it also responded to exogenous, as well as endogenous, cell wall stress. Surprisingly, vancomycin did not induce promoter activity in any of the strains except the Δ*tipN* Δ*chvT* double mutant (Fig 6D).

**Fig 6.**
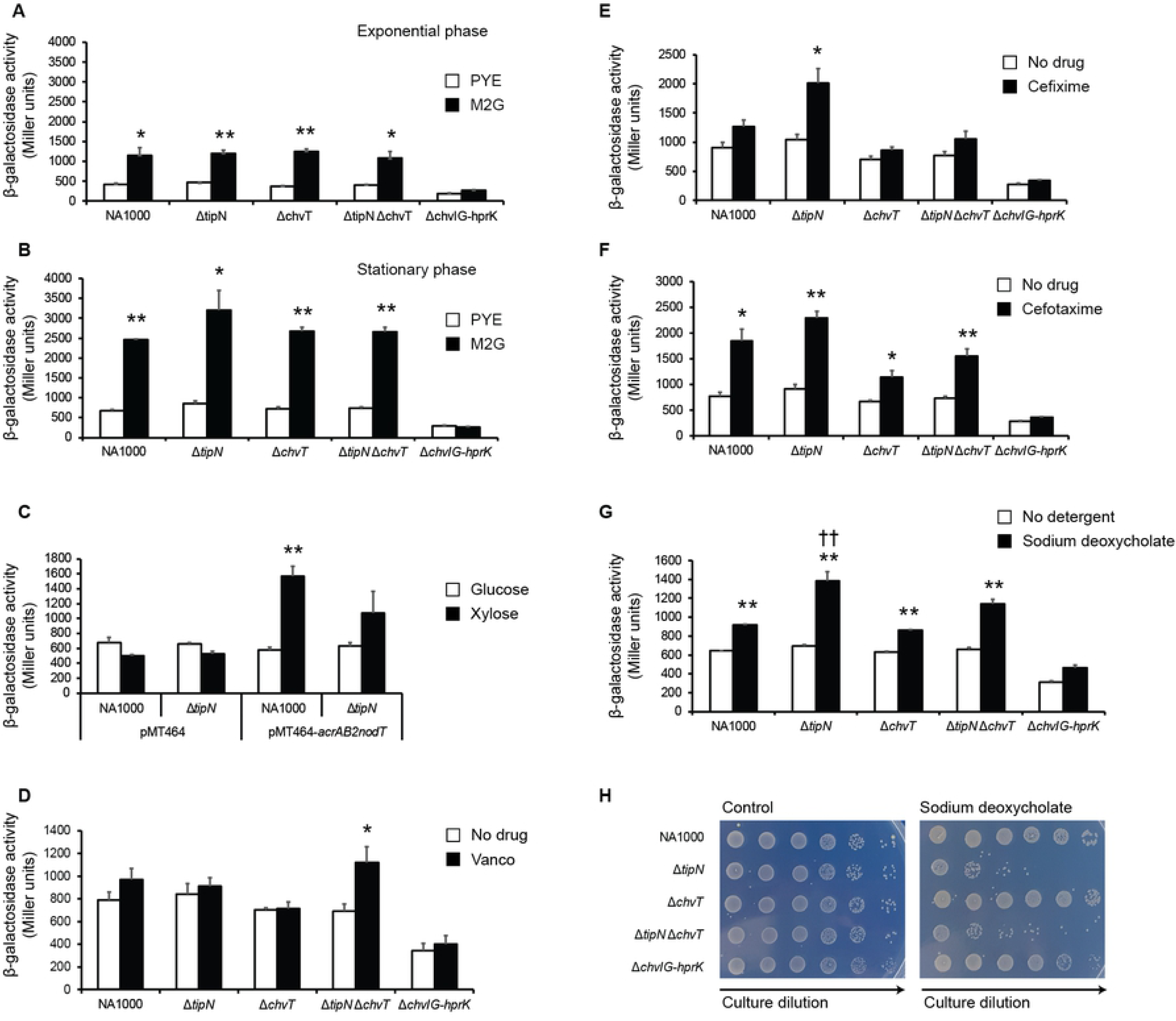
The ChvIG-ChvR-ChvT signalling pathway responds to cell wall stress in *Caulobacter*. (A) Promoter activity of P*_chvR_*in WT, Δ*tipN*, Δ*chvT*, Δ*tipN* Δ*chvT* and Δ*chvIG-hprK* in exponential phase (3hr growth after inoculation) in PYE or M2G media. (B) Promoter activity of P*_chvR_* in WT, Δ*tipN*, Δ*chvT*, Δ*tipN* Δ*chvT* and Δ*chvIG-hprK* in stationary phase (24hr growth after inoculation) in PYE or M2G media. These experiments were performed independently of those measured in (A) and are not derived from repeat measurements of the same cultures. (C) Promoter activity of P*_chvR_*in WT and Δ*tipN*, containing overexpression plasmid pMT464-*acrAB2nodT* or empty vector pMT464, treated with 0.3% xylose or 0.2% glucose for 24hr. (D) Promoter activity of P*_chvR_* in WT, Δ*tipN*, Δ*chvT*, Δ*tipN* Δ*chvT* and Δ*chvIG-hprK* with or without 15 μg/ml Vanco for 24hr. (E) Promoter activity of P*_chvR_* in WT, Δ*tipN*, Δ*chvT*, Δ*tipN* Δ*chvT* and Δ*chvIG-hprK* with or without 5 μg/ml cefixime (= 10 μM) for 24hr. (F) Promoter activity of P*_chvR_* in WT, Δ*tipN*, Δ*chvT*, Δ*tipN* Δ*chvT* and Δ*chvIG-hprK* with or without 5 μg/ml cefotaxime (= 10 μM) for 24hr. (G) Promoter activity of P*_chvR_* in WT, Δ*tipN*, Δ*chvT*, Δ*tipN* Δ*chvT* and Δ*chvIG-hprK* with or without 0.6 mg/ml sodium deoxycholate for 24hr. (H) Viability assay of WT, Δ*tipN*, Δ*chvT*, Δ*tipN* Δ*chvT* and Δ*chvIG-hprK* with or without 0.6 mg/ml sodium deoxycholate for 24hr under the same conditions as in (G). Images are representative of 3 independent experiments. Statistical significance is denoted by: * for p < 0.05 and ** for p < 0.01 in within-strain comparisons (all panels, control vs. test condition) and †† for p < 0.01 for between-strain comparison under the same experimental condition (WT vs. Δ*tipN* both with sodium deoxycholate, Fig 6G only).

However, the cephalosporin antibiotics did induce promoter activity in a strain-dependent manner. Cefixime only provoked increased P*_chvR_* activity in the Δ*tipN* strain (Fig 6E). Cefotaxime increased its activity in all the strains, although the magnitude of the increase was less in the Δ*chvT* and Δ*tipN* Δ*chvT* mutants compared to the WT and Δ*tipN* strains (Fig 6F). This increase was always ChvIG-dependent, as it was never seen in the Δ*chvIG-hprK* strain. Finally, we investigated the effect of inducing non-specific cell envelope stress on P*_chvR_* activity with sodium deoxycholate, a detergent which destabilises the outer membrane [16]. This condition increased P*_chvR_* activity in all strains, with a significantly larger increase seen in the Δ*tipN* strain compared to WT (Fig 6G). This increased activity correlated with a loss of cell viability in the Δ*tipN* strain compared to WT cells (Fig 6H). In sum, our data argue in favor of a novel structural role for TipN in stabilising the cell envelope at the weak points of poles and division plane, in addition to its developmental roles in cellular differentiation and replication, and show that the ChvIG-dependent signalling pathway in *Caulobacter* has been co-opted as a cell envelope stress sensor.

## Discussion

In the present study we have demonstrated that the loss of TipN in *Caulobacter* is associated with sensitivity to some types of cell envelope perturbation, either by overload of cell envelope-located multidrug efflux pumps (surprisingly, since these are otherwise beneficial to the cell), destabilisation of the outer membrane by detergents or exposure to antibiotics that target the cell envelope. This work also explains our previous observation that Δ*tipN* cells treated with the efflux inhibitor PAβN grew even worse than vehicle-treated control cultures [11], as this compound is known to have non-specific destabilising effects on the outer membrane as well as blocking efflux [29]. Since the subcellular location of TipN is restricted for the majority of the cell cycle to the new pole of the cell or to the division plane, it follows that if TipN is stabilising the cell envelope in some way, it is doing so at these places specifically. TipN co-localises with the Tol-Pal complex of *Caulobacter*, which is enriched at the new pole and division plane of the cells and is important for maintenance of cell envelope integrity during growth and division and for correct localisation of TipN during the cell cycle. In depletion strains lacking TolA, TolB or Pal, the poles and division plane frequently showed bleb formation caused by envelope layer separation [9]. In this study (and previously) we did not observe similar blebbing during normal growth of Δ*tipN* cells, but we did when the cells were exposed to Vanco. Therefore, if TipN is part of a Tol-Pal-TipN complex needed for full stabilisation of the cell envelope layers, TipN’s role in it seems to be required under stress conditions only. The incorporation of TipN into such a complex is probably through an interaction with the TolA protein, most likely through its inner membrane helices. TolA and TipN are both inner membrane proteins, but while TolA is anchored to the inner membrane by a single transmembrane (TM) helix with the rest of the protein in the periplasm, TipN has two inner membrane TM helices connected by a short (approximately 15 amino acids) periplasmic linker and the remainder of the protein is in the cytoplasm [5, 6]. It is also possible that the need for TipN in order to maintain cell envelope integrity during cell wall stress is not due to a direct structural role for TipN in the Tol-Pal complex, but a requirement for TipN to be present in order to execute the last stages of cytokinesis efficiently when the cell is under stress. Although TipN is dispensable for cell division in normal conditions, since cells lacking TipN are only moderately filamentous, TipN has been shown by bacterial 2-hybrid assay to interact with the late divisome component FtsN [30] and the bulging and filamentation phenotypes of Δ*tipN* (but not WT) cells that we observed upon AcrAB2-NodT efflux pump overexpression are consistent with this hypothesis. It is unlikely that TipN is needed for recruitment of divisome proteins, as it has been shown that TipN arrives at the division plane after the rest of the divisome has been assembled and the Tol-Pal complex has already arrived there [31]. However, our current data could also support a model where TipN is required for stabilisation of the divisome and in its absence the septal peptidoglycan biosynthetic machinery is more easily accessed or compromised by Vanco, leading to blebbing and lysis, instead of (or in addition to) a structural requirement for TipN for cell envelope integrity.

It was unexpected to see that the outer membrane-disrupting detergent sodium deoxycholate was the only factor tested here that showed both reduced viability of Δ*tipN* cells and a corresponding increase in *chvR* promoter activity, relative to WT. These findings suggested that this detergent causes a greater degree of stress to Δ*tipN* than to WT cells. However, TipN does not make any direct contact with the outer membrane in WT cells but is located in the inner membrane and cytoplasm. Therefore, if TipN is affecting the outer membrane it is probably doing it indirectly via an interaction with an outer membrane-located protein or complex. The Tol-Pal complex would be the most likely candidate for this, since the Pal protein is integrated in the outer membrane and contacts TolA and the rest of the complex in the periplasm. Our data are consistent with a model where TipN participates in the Tol-Pal complex’s function of holding the cell envelope layers together at the cell division site and that in the absence of TipN, the layers lose contact with each other more easily, which is imperceptible under normal conditions but leads to increased sensitivity to detergent acting at the outer membrane, efflux pump overexpression overloading the periplasmic space or antibiotics which act on the peptidoglycan (Fig 7). An alternative explanation could be that TipN is needed for clustering of Tol-Pal complexes at the division plane to increase their local density and reinforce the cell envelope at the division site during membrane fission and remodelling of the peptidoglycan, for example by each TipN protein contacting one Tol-Pal complex through an interaction between the TipN and TolA inner membrane helices and tethering them together through interactions between the TipN cytoplasmic coiled-coil domains. In the absence of TipN, the Tol-Pal complexes would be less densely concentrated at the division plane, making this site more fragile to external stress. Imaging experiments, eventually combined with construction of nested deletion of TipN inner membrane or cytoplasmic domains, would be required to test these hypotheses.

**Fig 7.**
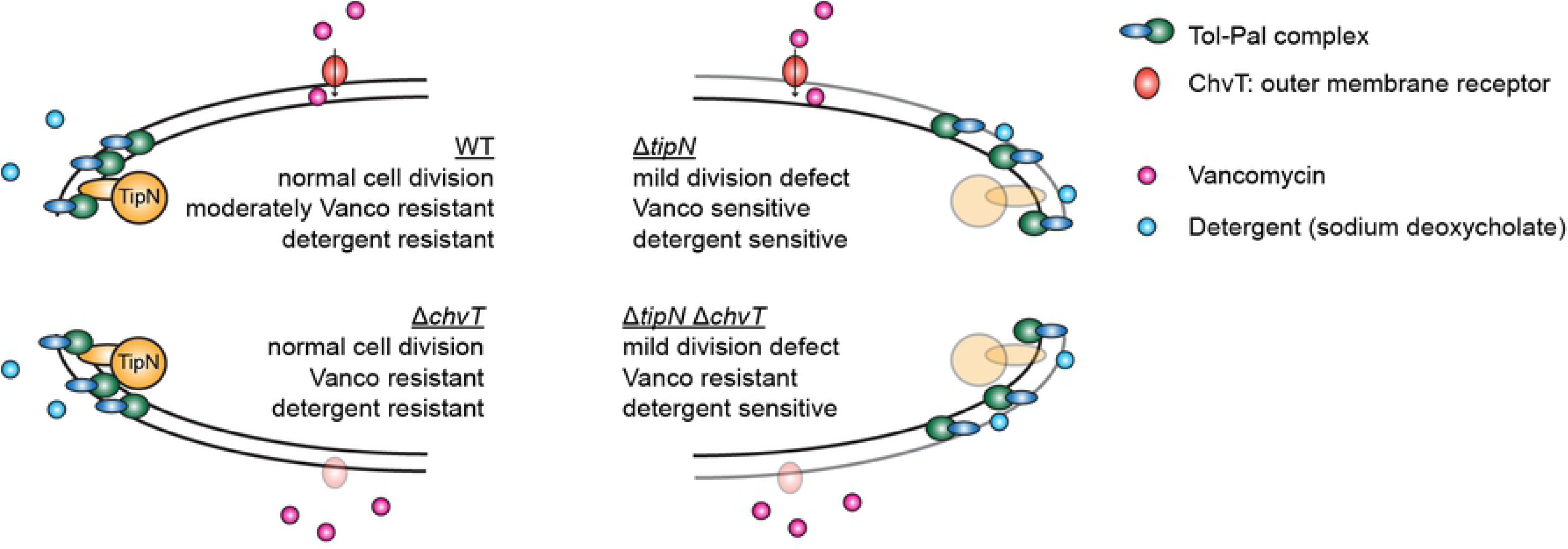
Loss of cell pole stabilisation by TipN sensitises cells to vancomycin-and detergent-induced cell wall stress. Summary graphical model of the relative effects of loss of TipN and/or ChvT on sensitivity to vancomycin and sodium deoxycholate in *Caulobacter crescentus*. The physical interaction between TipN and the Tol-Pal complex is known from previous work [9] while the different density of Tol-Pal complexes in WT vs. Δ*tipN* is our current working hypothesis but is not proven. For the purposes of this figure ChvT is illustrated at the lateral side of the cell, but it is likely to be distributed in multiple locations around the outer membrane [45].

Our forward genetic screen for suppressors of exogenous cell wall stress (Vanco) sensitivity in the Δ*tipN* strain revealed that growth in Vanco is improved by loss-of-function mutation of the TonB-dependent receptor *chvT*. Since this was observed in both WT and Δ*tipN* backgrounds, it seems unlinked to any of the intracellular functions of TipN and is more likely to arise from absence of ChvT making the cells’ outer membrane less permeable to the drug. However, this raises several questions about the normal function of ChvT. Since *Caulobacter* prefers oligotrophic aquatic habitats, it is well adapted for nutrient scavenging and encodes more than 60 TonB-dependent receptors in its genome, many of which have been seen to be upregulated during carbon or nitrogen starvation [17]. However, ChvT was not found among the TonB-dependent receptors that were upregulated during starvation conditions in that study, and our transcriptional reporter results show that it is constitutively produced, at least during exponential phase in rich media (Figure 5B). Indeed, given that transcription of the ChvT-downregulating small RNA ChvR is increased under starvation conditions (M2G media) and in stationary phase (Figure 6B and [22]) it appears that the reverse is true; ChvT should be produced more in rich media and less in nutrient-poor conditions, either minimal medium or in stationary phase, which are more similar to its natural growth conditions in the environment. The observed vancomycin sensitivity of *Caulobacter* (since even WT *Caulobacter* is more sensitive to vancomycin than other Gram-negative bacteria [32]) might therefore be linked to the relatively nutrient-rich conditions under which it is grown in standard laboratory culture, which are not representative of natural growth. It is of interest in this context that three specific polymorphisms in the *chvT* gene have been associated with adaptation to laboratory culture conditions at the expense of stationary phase survival [33]. These polymorphisms are missense mutations and therefore do not necessarily cause loss-of-function, but it is still unknown what either the polymorphisms, or the ChvT gene product itself, do. Based on homology to other TonB-dependent receptors, we would predict that ChvT is a gated channel that requires energy to open and bring in the substrate, not a pore that allows free diffusion, and that it should have some degree of substrate specificity. It could be speculated, based on the association of *chvT* with growth in and adaptation to nutrient-rich conditions, that it has specificity for peptide-based substrates and that Vanco, being a glycopeptide antibiotic, is toxic because it is similar enough to the natural substrate of ChvT that it is taken up by accident. If this were the case it would also explain why *chvT* deletion protected the cells so much more strongly against Vanco than it did against cefixime and cefotaxime, despite these antibiotics all sharing the same target.

We have shown, using a transcriptional reporter for the promoter of the ChvR sRNA, that it can be transcriptionally activated by cell wall-targeting antibiotics (in a strain-dependent manner) and by non-specific destabilisation of the outer membrane by the detergent sodium deoxycholate, in addition to the known stimulus of M2G minimal media (Figure 6). Since the only known transcriptional activator of ChvR is the two-component system ChvIG, and our results show that the increases in ChvR promoter activity in response to all stimuli were dependent on ChvIG, our data support a model where the sensor kinase ChvG senses cell envelope stress and activates the response regulator ChvI to induce transcription of its target genes, including *chvR*, which would result in post-transcriptional downregulation of ChvT. However, although *chvT* is the only target of ChvR, *chvR* is highly unlikely to be the only target of ChvI. Other genes of the ChvI regulon may be more important for cell wall stress tolerance, while post-transcriptional downregulation of ChvT could be a minor aspect of the response serving to reduce the permeability of the outer membrane to exogenous sources of cell wall stress that can use ChvT as their route of entry. The ChvI regulon has been defined in *Sinorhizobium meliloti* and contains many genes involved in symbiosis with the host plant cells, notably including genes that regulate cell envelope integrity [34]. Defining the *Caulobacter* ChvI regulon would clarify to what extent ChvI-dependent gene regulation patterns are shared or divergent between symbiotic and free-living alpha-proteobacteria and whether maintenance of cell wall integrity has taken over as the major role of ChvIG signalling in *Caulobacter*. Many responses to external stress factors in *Caulobacter* are mediated by ECF-family σ factors [35–37] and it will be intriguing to discover how ChvIG-mediated gene expression pathways are integrated with σ factor dependent regulation. The question remains open how the sensor kinase ChvG of *Caulobacter* can sense and integrate such different classes of signals such as nutrient starvation, acid stress and both exogenous and endogenous sources of cell wall stress. However, the orthologous sensor kinase ExoS of *S. meliloti* is influenced by the periplasmic regulatory factor ExoR which affects ChvI-dependent gene expression through physical interaction with ExoS [38]. Similar factors may also contribute to ChvIG signalling activation in *Caulobacter*, perhaps as dedicated transducers of cell wall stress signals to ChvG.

While ChvG appears to be a sensor of certain types of external stress (acid, nutritional, detergent and certain antibiotics) it is currently unclear how it is sensing them. It was unexpected, in this context, that we did not observe a strict correlation between stress factors to which Δ*tipN* cells were sensitised relative to WT and which loss of ChvT protected against, nor between factors which loss of ChvT protected against and ones which induced P*_chvR_* activity. The variety of stress stimuli which result in P*_chvR_* activation suggest that ChvG is able to sense stress that occurs both in the periplasm and at the outer membrane. However, although our data are consistent with ChvT taking up vancomycin from the external environment into the periplasm, based on the results of growth in vancomycin for WT compared to Δ*chvT* strains, we observed no correlation between P*_chvR_* activity and ChvT presence or absence in our promoter activity assays. Therefore, the presence or absence of ChvT does not appear to exert any feedback on ChvG sensor kinase activity, indicating that the ability of vancomycin to access the periplasm does not directly stimulate ChvG signaling. This would suggest that ChvG is primarily sensing at the outer membrane surface and that the induction of P*_chvR_*activity in WT cells on sodium deoxycholate exposure is a direct result of outer membrane destabilisation, while induction of its activity on overexpression of *acrAB2nodT* from the xylose inducible promoter, or on exposure to peptidoglycan-attacking antibiotics, would be indirect. Further experiments would be required to investigate whether this is the case and to investigate the reason why the two similar cephalosporin antibiotics cefixime and cefotaxime show moderately different profiles of P*_chvR_* activity between strains when they are otherwise similar in molecular structure and activity.

Free-living bacteria depend on the effectiveness of their signal transduction systems to communicate the presence of external stress factors to the genetic regulatory machinery and to alter gene expression, whether at the transcriptional or post-transcriptional level, in order to adapt and overcome them. We present evidence here that the two-component system ChvIG, exclusively associated until now with host-associated (pathogenic or symbiotic) alpha-proteobacteria, is also highly important for oligotrophic free-living bacteria as the signal transduction mechanism for alerting the bacteria to cell envelope stress. We further show that the system can modulate its activity in the event of altered intrinsic sensitivity to such stresses, as in the case of Δ*tipN* mutant cells which have higher levels of ChvIG-dependent gene activation during detergent-induced outer membrane stress, to which they are more sensitive than WT cells. TipN was already known to be a multifunctional “hub” protein required for correct asymmetric differentiation and for chromosome segregation in *Caulobacter*, and we now add another role to its repertoire, of maintaining polar cell envelope integrity. The implications of the findings presented here are twofold; first that the two-component system ChvIG may influence gene regulation in response to cell wall targeting antibiotics, such as might be used against pathogenic species of alpha-proteobacteria, and secondly that resistance to such antibiotics is dependent on factors such as TipN that maintain the integrity of the cell envelope at the otherwise vulnerable points of the cell poles and division plane. The possibility follows that polarly localised factors in other rod-shaped bacteria, not limited to the alpha-proteobacteria, may also play similar cell envelope stabilising roles and similarly function as antibiotic resistance determinants (and hold potential as drug targets).

## Materials and Methods

### General growth conditions

*Caulobacter crescentus* and *E. coli* strains were routinely grown in PYE at 30°C and LB at 37°C, respectively. Antibiotics for plasmid-borne selectable markers were used at the following concentrations: tetracycline at 1 μg/ml for *Caulobacter* and 10 μg/ml for *E. coli*, kanamycin at 20 μg/ml (solid media) or 5 μg/ml (liquid media) for *Caulobacter* and 20 μg/ml for *E. coli* and gentamicin at 1 μg/ml for *Caulobacter* and 10 μg/ml for *E. coli*. Preparation of M2G medium, electroporation, conjugations from *E. coli* to *Caulobacter* and generalized transduction with ΦCr30 bacteriophage were performed as described [39]. Stock solutions of vancomycin, cefixime and cefotaxime (sodium salt) were prepared at 50 mg/ml in water, 25 mg/ml in DMSO and 50 mg/ml in water, respectively. Stock solution of sodium deoxycholate was prepared at 60 mg/ml in water.

### Strain construction

Details of plasmid construction methods are in S1 Text and primer sequences and plasmid descriptions are in S1 Table and S2 Table respectively. High copy plasmids (pMT335, pMT464 [40] and their derivatives) were transferred into *Caulobacter* strains by electroporation while low copy plasmids and suicide plasmids (plac290 and pNPTS138 derivatives, respectively) were transferred by conjugation from *E. coli* S17-1 λ*pir* [41]. High and medium-copy plasmids (pET28a and pSRK-Km [42] derivatives) were transformed into *E. coli* strains by electroporation. After integration of suicide vectors by recombination through the first homology driver, secondary recombination events were induced by sucrose counterselection and mutants carrying resulting in-frame deletions were screened for by PCR. Strain numbers and genotypes are listed in S3 Table.

### End-point growth assays

To measure relative growth in liquid culture with antibiotics or during efflux pump gene overexpression, *Caulobacter* strains were grown to stationary phase, culture density (OD600) was measured and the starter cultures were diluted to a final density of OD600 = 0.001 (equivalent to 10^6^ cells/ml) into 5 ml PYE containing the antibiotic to be tested along with other antibiotic selection markers for plasmids. Where appropriate, 50 μM vanillate to induce gene expression from pMT335 derivatives, and 0.3% xylose to induce or 0.2% glucose to repress gene expression from pMT464 derivatives, was added. Cultures were grown for 20hr at 30 °C (to early stationary phase) and the OD600 measured. Data are expressed throughout as a percentage of the OD600 of a control culture of each strain, including plasmid selection markers where necessary but without any other supplement. All graphs show mean ± SEM from three independent experiments.

### Kinetic growth assays

Growth kinetics were measured in a temperature-controlled Synergy H1 multimode plate reader (Biotek) at OD600 during 48 hours incubation at 30°C. Starter cultures of *Caulobacter* strains were grown to stationary phase in PYE containing appropriate antibiotics for plasmid selection if necessary, then diluted and normalised to OD600 = 0.1. Vancomycin, xylose or glucose was supplemented as specified in the Supporting Information captions and 200 μl of the diluted cultures were transferred into duplicate wells of a clear flat-bottomed 96 well plate. OD600 was read once per hour with plate agitation for 15 seconds before measurement. All graphs show mean and standard deviation for all data points of three independent experiments.

### AcrA protein purification and western blotting

His6-AcrA was overexpressed by addition of 1 mM IPTG from pET28-*acrA* in *E. coli* Rosetta and purified under native conditions by Ni-NTA chelate chromatography. Purified His6-AcrA protein was used to immunize rabbits (Josman LLC, USA) for antiserum production. Cell extracts were prepared from mid-exponential phase cultures normalized to culture density, electrophoresed in 15% SDS-polyacrylamide gels and the proteins transferred to PVDF membranes. The anti-AcrA antiserum was diluted to 1/50’000 and detected by a horseradish peroxidase-coupled secondary antibody with chemiluminescent substrate.

### Microscopy

Cultures for imaging were immobilized on 1% agarose (in water) slides. DIC images were obtained with an Axio Imager M2 microscope equipped with a 100x/1.46 oil objective and a CoolSnap HQ^2^ camera. Phase-contrast images were obtained with an inverted Olympus IX83 microscope with a Photometrics Prime sCOMS camera. All images shown are representative of three independent experiments.

### Chemical library screening

The Prestwick Chemical Library was dispensed into 96 well plates so as to have 2 wells (in different plates) per compound. Each plate contained 8 wells negative control (DMSO), 8 wells positive control (chloramphenicol) and 80 wells of library compounds. Saturated overnight cultures of WT cells were diluted to an OD600 of 0.001 in 2 x 800 ml PYE and Nal was added to one at a final concentration of 20 μg/ml. The chemical library plates were allowed to come to room temperature, then centrifuged at 2000 rpm for 4 minutes. 200 µl starter culture was seeded into each well, lids were replaced, and plates incubated at 30°C for 48hr at which point cells in the negative control wells had reached stationary phase. Optical density (630 nm) was read and the readings compiled into a single Excel spreadsheet per screen. The person performing this experiment was blinded to the position of the library compounds in the plates. For data analysis, a score was calculated for all compounds normalised to the mean of the positive control wells as the upper limit (=1) and the mean of the negative control wells as the lower limit (=0). Nal-dependent hits were considered as those whose score fell outside the boundary of 3 standard deviations from the mean of the negative controls in the screen that was performed in the presence of Nal, but not in the control screen.

### Transposon mutant library generation, selection of vancomycin resistant mutants and mapping of transposon insertions

The transposon mutant pooled library was created by electroporation of the transposon-bearing plasmid pMR2xT7 into *Caulobacter crescentus* Δ*tipN* followed by selection on gentamicin. After incubation at 30°C for 4 days, approximately 20’000 individual colonies were collected into a final volume of 13 ml PYE and stored at −80°C in 1 ml aliquots containing 10% DMSO. For selection of Vanco resistant clones by enrichment culture, three overnight cultures were set up from separate frozen aliquots. Upon reaching stationary phase they were diluted 1/1000 into PYE containing 15 μg/ml Vanco and incubated at 30°C for 24 hours. 100 μl aliquots were plated on solid PYE media containing 1 μg/ml gentamicin and 15 μg/ml vancomycin and incubated at 30°C for 3 days. Single colonies were picked and re-streaked twice to new PYE plates with these concentrations of gentamicin and Vanco to ensure stability of the Vanco resistance. Cultures of these re-streaked clones were used as template for transposon mapping by 2-step arbitrary PCR as in [43]. Briefly, the first arbitrary PCR reaction was performed in a total volume of 25 μl containing 2 μl overnight culture of the transposon mutant strains as template, 1 ng/μl of each of primers pMar2xT7_Arb1_A and pMar2xT7_Arb1_B, 2.5 μM dNTPs, 10% DMSO and 1.25 U Taq polymerase. 5 μl of this first PCR reaction was used as template for the second arbitrary PCR reaction which contained identical dNTP, DMSO and Taq concentrations and 1 ng/μl of each of primers pMar2xT7_Arb2_A and pMar2xT7_Arb2_B, in 25 μl. PCR products from these reactions were purified by agarose gel electrophoresis and sequenced with the nested sequencing primer pMar2xT7_Arb3_A. The three isolated clones with mutations in or near *chvT* (1h15, 2v15 and 3l15) had the transposon insertions backcrossed into the Δ*tipN* strain by ΦCr30 bacteriophage transduction to confirm dependence of Vanco resistance on the transposon insertions.

### β-galactosidase assays

β-galactosidase assays on strains carrying plasmid-borne transcriptional fusions of test promoters to *lacZ* were performed by the method of Miller [44] at 30°C on cells exposed to antibiotics or other inducers (as described in main text or figure legends) for three hours for exponential phase or 24 hours for stationary phase, from three independent biological replicates. Stationary phase cultures were diluted ¼ to obtain an OD660 measurement in the range of 0.1 – 0.4 before performing the assay. All graphs show mean ± SEM from three independent experiments unless otherwise stated in the figure legends.

### Sodium deoxycholate resistance assay

Sodium deoxycholate resistance was assayed as described in [16]. Cultures of strains to be tested were diluted from saturated overnight cultures to an OD600 of 0.2 in PYE with and without sodium deoxycholate at 0.6 mg/ml and incubated at 30°C for 24 hours. Culture density was measured, normalised to the OD600 of the least dense culture, serially diluted (tenfold) in PYE to 10^-6^ and 5 μl of the dilutions spotted onto PYE plates. Plates were imaged after 48hr growth at 30°C. Three independent biological replicates were performed.

### Statistical analysis

Statistical significance was analysed by paired-sample 2-tailed Student’s T test for within-strain (xylose to glucose) comparisons (Figs 1 and 2), and by non-paired equal variance 2-tailed Student’s T test for between-strain comparisons of growth (Fig 5) or of β-galactosidase activity (Fig 6). * signifies p < 0.05 and ** signifies p < 0.01 throughout.

## Supporting Information captions

**S1 Fig. Shorter exposure images of the anti-AcrA immunoblots of Fig 1B and 1C.**

**S2 Fig. AcrAB2-NodT overexpression causes Δ*tipN* cultures to prematurely enter stationary phase.** Growth curves of WT and Δ*tipN* strains with empty vector control (pMT464) or AcrAB2-NodT overexpression plasmid (+ABN) grown either in non-supplemented PYE or in PYE with 0.3% xylose to induce overexpression, in 96-well plate format at 30°C for 48 hours. Control cultures containing 0.2% glucose (represses the xylose-inducible promoter) were also performed in all replicates of the experiment and always grew better (to a higher stationary phase OD600) than cultures in PYE alone.

**S3 Fig. High throughput chemical-genetic screening suggests Δ*tipN* cells are also sensitive to the cephalosporin antibiotics cefixime and cefotaxime.** (A) Scatter plot of the same screening data shown in Figure 4A, of WT *Caulobacter crescentus* (NA1000) grown in the presence of 20 μg/ml Nal in addition to the Prestwick Chemical Library compounds, with the points corresponding to cefixime wells (purple) and cefotaxime wells (turquoise) indicated. (B) Scatter plot of Δ*tipN* cells screened against the Prestwick Chemical Library with the points corresponding to cefixime wells (purple) and cefotaxime wells (turquoise) indicated. This screening experiment was previously published (Kirkpatrick et al., 2016); its Z’ score was 0.894.

**S4 Fig. *chvT* deletion does not improve the growth of Δ*tipN* in Nal or during overexpression of the AcrAB2-NodT efflux pump.** (A) Growth assay of Δ*tipN*, Δ*tipN* Δ*chvT*, Δ*tipN chvT::2v15* and Δ*tipN chvT::3l15* treated with 20 μg/ml Nal, 15 μg/ml Vanco or both antibiotics together for 20hr before culture density measurement. Data are expressed as % of OD600 of control cultures of each strain in untreated PYE. (B) Growth assay of Δ*tipN* Δ*chvT* cells with empty vector pMT464 and the series of pMT464 overexpression plasmids containing *acrAB2nodT* component genes separately or together, treated with 0.3% xylose or 0.2% glucose for 20hr before culture density measurement. Data are expressed as % of OD600 of control cultures of each strain in untreated PYE.

**S5 Fig. Vanco causes premature entry of Δ*tipN* and Δ*acrAB2nodT* strains into stationary phase.** Growth curves of WT, Δ*tipN*, Δ*acrAB2nodT*, Δ*tipN* Δ*acrAB2nodT*, Δ*chvT* and Δ*tipN* Δ*chvT* in PYE with and without 15 μg/ml Vanco, in 96-well plate format at 30°C for 48 hours.

**S6 Fig. *chvT* deletion protects against cefixime and cefotaxime in addition to vancomycin, but *chvT* overexpression does not increase sensitivity to these antibiotics.** (A) Growth assay of WT, Δ*tipN*, Δ*chvT* and Δ*tipN* Δ*chvT* treated with cefixime at the indicated concentrations for 20 hours before culture density measurement. Data are expressed as percentage of OD600 of control cultures of each strain in untreated PYE. (B) Growth assay of WT and Δ*tipN* containing the overexpression plasmid pMT335-*chvT* or the pMT335 empty vector, treated with 50 μM vanillate and cefixime at the indicated concentrations for 20 hours before culture density measurement. Data are expressed as percentage of OD600 of control cultures of each strain in PYE containing 50 μM vanillate only. (C) Growth assay of WT, Δ*tipN*, Δ*chvT* and Δ*tipN* Δ*chvT* treated with cefotaxime at the indicated concentrations for 20 hours before culture density measurement. Data are expressed as percentage of OD600 of control cultures of each strain in untreated PYE. (D) Growth assay of WT and Δ*tipN* containing the overexpression plasmid pMT335-*chvT* or the pMT335 empty vector, treated with 50 μM vanillate and cefotaxime at the indicated concentrations for 20 hours before culture density measurement. Data are expressed as percentage of OD600 of control cultures of each strain in PYE containing 50 μM vanillate only. Statistical significance in panels A and C is denoted by: * for p < 0.05 and ** for p < 0.01 in WT to Δ*tipN* comparison, † for p < 0.05 in WT to Δ*chvT* comparison and § for p < 0.05 in Δ*tipN* to Δ*tipN* Δ*chvT* comparison.

**S1 Table. Primers used in this study.**

**S2 Table. Plasmids used in this study.**

**S3 Table. Bacterial strains used in this study.**

**S1 Text. Details of plasmid construction.**

